# Personalized connectivity-based network targeting model of TMS for treatment of psychiatric disorders: computational feasibility and reproducibility

**DOI:** 10.1101/2023.06.28.545400

**Authors:** Zhengcao Cao, Xiang Xiao, Cong Xie, Lijiang Wei, Yihong Yang, Chaozhe Zhu

## Abstract

Consider the limited clinical efficacy of transcranial magnetic stimulation (TMS) due to heterogeneity in treatment outcomes, the utilization of individual functional connectivity (FC) can enhance the prediction accuracy in the network targeting model. However, the low signal-to-noise ratio (SNR) of FC poses a challenge when utilizing individual resting-state FC (rsFC). To overcome this challenge, proposed solutions include increasing the scan duration and employing clustering approaches to enhance the stability of FC. In this study, we aimed to evaluate the stability of a personalized functional-based network targeting model in individuals with major depressive disorder (MDD) and schizophrenia with auditory verbal hallucinations (AVH). Using resting-state functional magnetic resonance imaging (rsfMRI) data from the Human Connectome Project (HCP), we assessed the model’s stability and employed longer scan durations (7 minutes, 14 minutes, 21 minutes, 28 minutes) and clustering methodologies to improve the precision of identifying optimal individual sites. Our findings demonstrate that a scan duration of 28 minutes and the utilization of the clustering approach lead to stable identification of individual sites, as evidenced by the intraindividual distance falling below the ∼1cm spatial resolution of TMS. These findings contribute to the understanding of individualized TMS targeting and have implications for improving treatment outcomes in psychiatric disorders.

## 1. Introduction

Transcranial magnetic stimulation (TMS) is a non-invasive neuromodulation technology with ∼1cm level spatial resolution. TMS has received FDR approval as a safe and effective therapy for MDD patients who do not respond to the behavioral or pharmacological treatment and has also proved its potential as novel treatment for other psychiatric disorders including schizophrenia. Though the general efficacy as is demonstrated for the TMS-based treatment, its clinical utility is limited by the heterogenous outcome on individual patients, even when their clinical conditions are similar.

Differences in the morphology and functional connectivity of individual brain may account for the heterogeneous outcome of TMS. Traditionally, TMS coil is set according to scalp landmarks, e.g. F3, anterior 5-cm from the MEP hop-spot for MDD or the mid-point of T3 and P3 for auditory verbal hallucinations (AVH) of schizophrenia. Such location-based targeting strategies, as well as that using more advanced neuronavigation technique, may oversimplify the physiological process how TMS generate the modulation effect on the human brain system. First, the E-filed distribution highly depends on the intracranial geometry of human brain (Thielscher et al., 2011). Thus, even when TMS coil is set in the identical spot of the patients’ scalp, the actual excited cortical area can vary greatly among different subject, or even on the same subject but with varied coil orientations. Second, the associative cortical areas that are commonly targeted for treating psychiatric disorders, e.g. the DLPFC for MDD and TPJ for AVH, exhibit the highest levels of interindividual variance in terms of structural morphology, neuronal function, and connection (Rajkowska and Goldman-Rakic, 1995; Fischl et al., 2008; Hill et al., 2010; Fox et al., 2013; Mueller et al., 2013; Mira-Dominguez et al., 2014; Finn et al., 2015; Doucet et al., 2019). As the result, varied network can be engaged in the effect field of the TMS stimulation, through the mono-/multi-synaptic connections to the local areas that directly receiving TMS stimulations, which is considered to account for the heterogeneous treatment efficacy of TMS.

On observation that the treatment efficacy is associated with the extend how the pathological network of given disorder is engaged in the stimulation network, our previous work (Cao et al., 2023) proposed the network targeting model (NTM) for guiding TMS coil placement for individual patients. Considering the reliability of the targeting result, the stimulation network of NTM was originally based on group-level FC.

The use of individual functional connectivity data has several advantages over normative or average connectomes (Klooster et al., 2022). First, a study compared group-based targeting with individualized targeting in transcranial magnetic stimulation (TMS) and found that individualized stimulation sites improved the reliability of TMS-evoked responses, particularly in highly variable task-positive networks, such as the Dorsal Attention Network (DAN) (Menardi et al., 2022). Second, studies comparing individual and normative connectomes have shown similar results in predicting clinical responses, but a trend towards better prediction was observed with individual data (Fox et al., 2013; Cash et al., 2020; Siddiqi et al., 2021; Kong et al., 2022).

A major challenge for incorporating individual FC into the NTA framework is in its relatively low SNR of rsfMRI. The low signal-to-noise ratio of rsfMRI in functional connectivity calculations can lead to inaccurate measurements of correlation values, as the weak brain signal compared to noise may overshadow the true underlying functional connectivity patterns (Birn, 2014; Mueller, 2015; Teeuw 2021). For this reason, it makes FC-based approaches unstable (Dubois and Adolphs, 2016) and give ambiguous guidance for setting the TMS coil (Ning et al., 2018).

Currently, of strategies have been proposed to reduce the spurious FC variance introduced by the data acquisition. One strategy to enhancing the stability of functional connectivity is to augment the number of data points or repetitions in resting-state functional magnetic resonance imaging . By extending the scan duration, a more comprehensive and stable evaluation of FC can be achieved, attributed to the reduction of noise (Birn 2013), increased statistical power, and capture of the temporal dynamics of brain activity (ref). Previous studies have demonstrated the beneficial impact of increased scan duration on the stability of individual rsFC (Finn et al., 2015; Gratton et al., 2018).

When FC is stable, it may indicate the presence of a robust pathway through which TMS exerts its effects on individuals. The establishment and stability of this pathway enable more precise and effective targeting of specific brain regions, which in turn contributes to more favorable treatment outcomes. Another category of strategy is to improve the stability by spatially averaging the rsFC map, or the ‘clustering’ approach which calculate the center-of-gravity of the largest cluster (Cash et al., 2021; Zhao et al., 2022). In MDD, the clustering approach has demonstrated its utility in reduce the within-subject instability while keeping the between subject variance for the optimized treatment site.

In the present study, our objective was to evaluate the stability of personalized functional-based network targeting model in individuals with major depressive disorder (MDD) and schizophrenia with auditory verbal hallucinations (AVH). We utilized two-day resting-state functional magnetic resonance imaging scans from the Human Connectome Project (HCP) dataset to assess the instability of the model, specifically focusing on the stimulation network, NTA map, and intraindividual distance. To address this stability challenge, we employed two strategies: longer scan time and the cluster methodology, aimed at improving the precision of identifying the optimal individual site. To ensure the generalizability of the model across different psychiatric disorders, we conducted stability validation in both MDD and schizophrenia with AVH by targeting MDD or schizophrenia with AVH pathological networks.

## 2. Methods

### 2.1. Overview

The aim of this study is to examine the variance of a personalized functional-based network targeting model using different rsfMRI scans and to reduce the variances through two strategies. In comparison to a network targeting model based on group-level rsFC, the personalized functional-based network targeting model utilizing individual rsFC can better capture interindividual differences (Supplementary Figure 1). Regarding the variance of individual functional connectivity, there are two sources of variance. The first is the desirable variance, including inter-individual differences in network organization and connectivity strength (Finn et al., 2015; Gratton et al., 2018). The second is the undesirable variance, including unwanted technical effects or the influence of rsfMRI nuisance variables (Birn, 2012; Birn et al., 2013; Bright and Murphy, 2015), which contribute to the variances observed in the stimulation network, NTA map, and optimal targets (Figure 1). In this study, we aim to assess these technical variances and propose two strategies to mitigate them: extending the duration of rsfMRI scans to enhance the signal-to-noise ratio of individual rsfMRI data and employing alternative searching methods to identify optimal targets.

**Figure 1.**
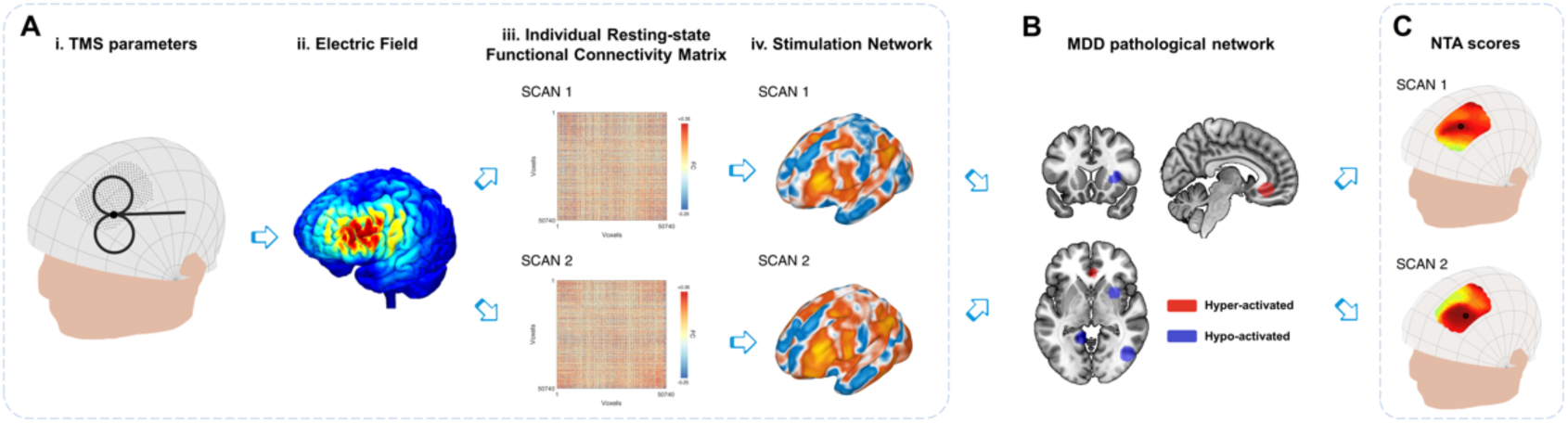
Variations in the personalized NTA model. **(A)** Stimulation network: TMS administered with specific combinations of parameters (i) from the search space results in an E-field (ii) that directly affects the local cortical region. The variances in individual resting-state functional connectivity from different scans (iii) contribute to the variations in the stimulation network (iv) within the stimulated cortical region. **(B)** Comparison with MDD pathological networks: The stimulation networks exhibiting spatial anti-correlation represent the MDD NTA score, which reflects the estimated efficacy of TMS (Cao et al., 2023). **(C)** For the entire search space, the NTA map is generated by considering all parameter combinations, with the optimal target site indicated by a black circle. The figure illustrates the variations in the NTA map and optimal target site.

### 2.2. Participants

A total of 134 participants [80 females, age 29.7 ± 3.5 years] were randomly selected from the Human Connectome Project (HCP) database (https://db.humanconnectome.org/). The rsfMRI data acquisition parameters in the database were TR=720ms, TE=33ms, flip angle=52°, FOV=208×180 mm², voxel size=2×2×2 mm³, and a multi-band factor of 8. The anatomical MRI volume size was 0.7×0.7×0.7 mm³. The anatomical MRI and rsfMRI data of the participants were used to construct E-field and rsFC separately.

Each participant underwent four fMRI scans on consecutive days. Two data acquisition sessions were conducted on each day, with each session comprising two 14-minute and 33-second runs (1200 volumes each) with right-to-left and left-to-right phase encodings. During scanning, participants were instructed to keep their eyes open and fixate on a projected bright cross-hair on a dark background.

### 2.3. rsfMRI data pre-processing

The rsfMRI data from the HCP dataset were preprocessed with the DPABI toolbox (Yan et al., 2016), which included the following steps: 1) elimination of the first ten time points; 2) correction for slice timing; 3) realignment of the functional image to correct for head motion; 4) regression of nuisance signals estimated from the signals of white matter, CSF and the mean global signal of gray matter (Fox et al., 2012); 5) 0.01∼0.1Hz band-pass filtering; 6) The functional images were co-registered to scalp-extracted anatomical images and then normalized into MNI space (3 mm × 3 mm × 3 mm) with the DARTEL algorithm (Ashburner, 2007) and 7) spatial smoothing (kernel FWHM 6 mm × 6 mm × 6 mm).

### 2.4. Compute personalized NTA for MDD and schizophrenia with AVH

#### 2.4.1. Search Space

In our study, for MDD, we utilized a cranial search space that covered a broad area around the left dorsolateral prefrontal cortex (DLPFC), consisting of 462 CPC positions and 20 mm radius spheres centered at BA9 [-36, 39, 43], BA46 [-44, 40, 29], Beam-F3 [-37, 26, 49], and 5cm [-41, 16, 54] (Lefaucheur et al., 2014; Xiao et al., 2018; Cash et al., 2021). The coordinates in the search space were *p_NZ_* ∈[0.15, 0.43] and *p_AL_*∈[0.27, 0.43], as shown in Supplementary Figure 2Ai. Similarly, for schizophrenia with AVH, we utilized a cranial search space consisting of 246 CPC positions and covering the left temporoparietal junction (TPJ) and left Wernicke Region, with 20 mm radius spheres centered at TPJ [-57, -49, 28] and L.Wernicke [-65, -41, 9] (Herwig et al., 2003; Hoffman et al., 2003, 2013). The coordinates in the search space were *p_NZ_*∈[0.52, 0.80] and *p_AL_*∈[0.10, 0.26], as shown in Supplementary Figure 2Bi. The CPC positions (*p_NZ_*, *p_AL_*) remained the same across the subjects, and the distance between two adjacent CPC positions on the individual head model was approximately 2.83 mm.

#### 2.4.2. Calculate individual network targeting accuracy map

##### 2.4.2.1. TMS coil placements

We used SimNIBS 3.2 (Thielscher et al., 2015; Saturnino et al., 2019) to segment T1 images of 134 participants and generate individual parameter spaces using their head surface nodes (Jiang et al., 2022). For MDD, a total of 462 coil placements (462 positions × 1 orientation) were used for calculation for each individual. The coil orientation was fixed at 45° from the midline, and the coil handle pointed backward (Fitzgerald et al., 2003; Thomson et al., 2013)(Supplementary Figure 2Aii). For schizophrenia with auditory verbal hallucinations (AVH), the handle direction was perpendicular to the TP3 line, which was around 23° from the midline measured within the Scalp Geometry-based Parameter (SGP) coordinate system (Jiang et al., 2022). The handle pointed backward (Paillère-Martinot et al., 2017)(Supplementary Figure 2Bii). A total of 246 coil placements (246 positions × 1 orientation) were set for each individual.

##### 2.4.2.2. Network targeting accuracy (NTA)

We created an electric field for an individual coil placement using SimNIBS 3.2 and assigned default isotropic tissue conductivities (Wagner et al., 2004). We selected the Magstim 70 mm figure-of-8 coil for electric field simulations (Thielscher and Kammer, 2004), following which we created individual E-field weights using the previous pipeline (Cao et al., 2023). The practical scans for clinical uses range from 6 to 8 minutes (Liu et al., 2017; Park et al., 2020; Kong et al., 2022), so we used half of right-to-left functional images of individuals (about 7 minutes) for the construction of individual rsFC. To determine individual rsFC, we masked the spatially normalized functional images of individuals in NMI space with a customized gray-matter mask consisting of 50740 voxels (Cao et al., 2023). We constructed a voxel-to-voxel correlation matrix (50740×50740) for each individual using Pearson’s correlation. Next, we built the TMS stimulation network of the coil placement using the individual E-field weights of the coil placement and individual voxel-to-voxel rsFC in MNI space. Finally, we determined the NTA by spatially anti-correlating the TMS stimulation network of the coil placement with the pathological networks derived from meta-analysis results for MDD or schizophrenia with AVH (Kühn and Gallinat, 2012; Gray et al., 2020; Cao et al., 2023).

##### 2.4.2.3. Individual NTA map

We computed NTAs for all coil placements targeting the pathological network in MDD (Supplementary Figure 2Ai & ii) and in schizophrenia with AVH (Supplementary Figure 2Bi & ii). The resulting NTA maps were displayed on individual head surfaces for MDD (Supplementary Figure 2Aiii) and schizophrenia with AVH (Supplementary Figure 2Biii).

##### 2.4.2.4. Individual scalp site

We used the “Classic” method (Cao et al., 2023) to identify the optimal scalp position for stimulation, determined by the maximum value of the individual NTA map. The three-dimensional coordinates of the optimal scalp position served as the individual stimulation site.

### 2.5. Evaluate the instability of personalized NTA model

We used three indices to evaluate the variance of the personalized NTA model, which included intrasession stimulation network similarity, intrasession NTA map similarity, and intraindividual distance.

Intrasession stimulation network similarity: To ensure the consistency of the stimulation network, it should be reliably determined by rsfMRI scans conducted on different days within the same individual. Therefore, the similarity between two separate stimulation networks obtained from the same individual should be maximized. The similarity was calculated as the correlation coefficient between the two stimulation networks.

Intrasession NTA map similarity: Similarly, the NTA maps should be consistently determined by rsfMRI scans performed on different days within the same individual. The similarity of two separate NTA maps obtained from the same individual should be maximum. The similarity was also calculated as the correlation coefficient between the two NTA maps.

Intraindividual distance: Intraindividual distance was used as an evaluation index to determine the optimal scalp site consistently from rsfMRI scans conducted on different days within the same individual. The distance between the optimal scalp sites calculated from two separate rsfMRI scans from the same individual should be minimum and less than the ∼1 cm spatial sensitivity of TMS (Deng et al., 2013), which is conventionally computed between two cortical sites (Ning et al., 2018; Cash et al., 2021; Du et al., 2022) due to TMS’s spatial sensitivity being described on the cortex (Deng et al., 2013). First, we projected the individual scalp coordinate onto the cortex by finding the closest cortical site along the normal vector. Then, we transformed the cortical coordinate into MNI space using the subject2mni_coords function of SimNIBS 3.2 package (Saturnino et al., 2019). Finally, the intraindividual distance was determined as the distance between the cortical coordinates of two separate scans conducted within the same individual.

### 2.6. Evaluate the feasibility of strategies for improving stability of personalized NTA model

Two strategies were implemented to enhance the stability of the personalized NTA model, namely the extension of rsfMRI scan durations and the cluster method. In this analysis, we present both strategies and the corresponding evaluation indexes.

i. Extend rsfMRI scan durations. We utilized longer rsfMRI scan durations. Specifically, we temporally concatenated two 14-minute 33-second runs per day to result in 28 minutes of data (Ning et al., 2018; Cash et al., 2021). We then divided the 28 minutes of data into four different scan durations: 7 minutes, 14 minutes, 21 minutes, and 28 minutes. Expect 7 minutes scan duration, we utilized other scan durations in the personalized NTA model to calculate intrasession stimulation network similarity, intrasession NTA map similarity, and intraindividual distance. This allowed us to observe the effect of longer rsfMRI scan durations on the stability of personalized NTA model.

ii. “Cluster” method (Cash et al., 2021): We utilized alternative searching methods to identify optimal targets. Since the NTA model starts on the scalp surface, we modified the “cluster” method from Cash (Cash et al., 2021), which starts on the cortical surface. We identified contiguous clusters from the individual NTA map (Figure 4Aii) and defined the center-of-gravity of the largest cluster as the target CPC position. This position’s three-dimensional coordinates were then used to determine the individual stimulation scalp site (Figure 4Aii). To define clusters, we used the top x% of NTA values, with the threshold ranging from 0.5% to 70%. We also limited clusters to the supra-threshold CPC points using 6 neighborhoods on a 2-D plane. Apart from intraindividual distance, we also employed interindividual and ratio of interindividual-to-intraindividual distance to evaluate the feasibility of the personalized NTA model. The interindividual distance was used to ensure that the personalized targeting results did not converge to a fixed scalp site, and that the optimal scalp site retained spatial diversity between individuals. Given that individuals have different head sizes, the interindividual distance was measured in MNI space (Balderston et al., 2020), which required projecting the individual scalp coordinate onto the cortex and transforming it into MNI space. The ratio of interindividual-to-intraindividual distance was used to ensure that the personalization methodologies maintained a smaller intraindividual distance and a larger interindividual distance, resulting in a higher ratio.

## 3. Results

### 3.1. Observe the variance of personalized NTA model

When we incorporated individual rsFC into the network targeting model, we observed significant variations in the stimulation network, NTA map, and the optimal target. For instance, Figure 2A and Figure 2B display the similarity and dissimilarity of the stimulation networks obtained from two 7-minute scans targeting points F3 and TP3. As shown in the middle row of Figure 2A (i.e., Sub ID 783462), the stimulation network obtained from two scans of the same target was relatively similar. However, in the bottom row of Figure 2A (i.e., Sub ID 189450), two scans of the same target resulted in different stimulation networks. We computed the intrasession stimulation network similarity for F3 and TP3, and the correlation coefficients were 0.569 and 0.600, respectively (Table 1). For all coil placements in the search space in targeting MDD and schizophrenia with AVH pathological networks, the averaged intrasession stimulation network similarity was under 0.6.

**Figure 2.**
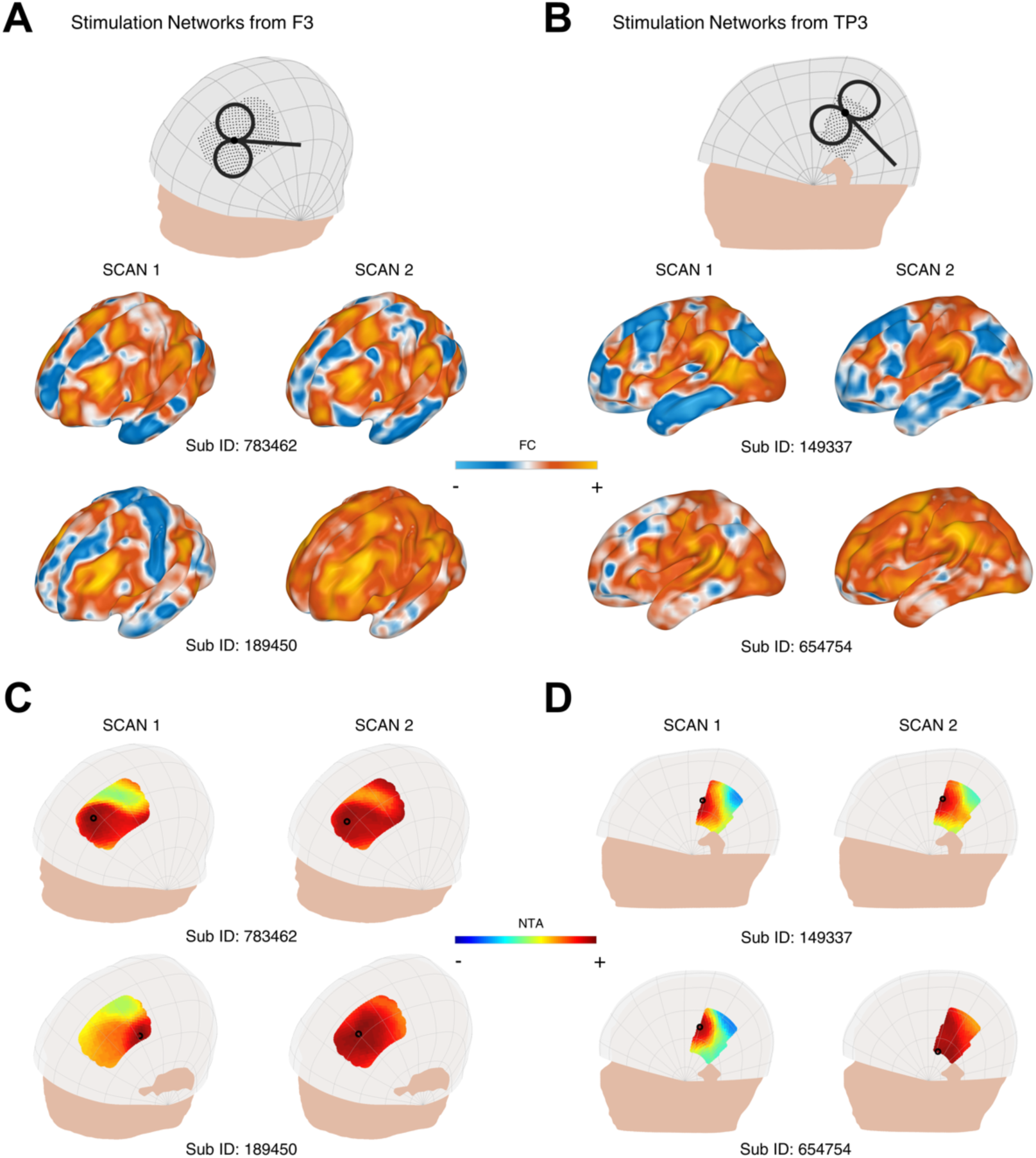
Variability of the personalized NTA model across different rsfMRI scans for targeting the pathological networks of MDD and schizophrenia with AVH. **(A)** shows the stimulation networks obtained from the F3 region, while **(B)** displays the stimulation networks from TP3 (the midpoint between T3 and P3). In the middle row of Figure 2A (Sub ID 783462), the stimulation networks derived from two scans of the same target exhibit relatively high similarity. However, in the bottom row of Figure 2A (Sub ID 189450), the same target produces different stimulation networks. NTA maps of representative individuals targeting the MDD pathological network are presented in **(C),** and the schizophrenia with AVH pathological network is shown in **(D)**. The individual scalp sites (indicated by black circles) were determined using the classic method. The individuals depicted in the first columns of Figure 2C show a wide range of target sites. In the middle row of Figure 2C (Sub ID 783462), the target sites remain consistent across individuals on separate days, whereas in the bottom row of Figure 2C (Sub ID 189450), the target sites vary among individuals on separate days.

**Table 1.** The variance of personalized NTA model.

|  | MDD | SZ with AVH |
| --- | --- | --- |
| Intrasession Stimulation Network Similarity of F3 [r] | $0.569 \pm 0.012$ | -- |
| Intrasession Stimulation Network Similarity of TP3 [r] | -- | $0.600 \pm 0.011$ |
| Intrasession Stimulation Network Similarity [r] | $0.553 \pm 0.009$ | $0.580 \pm 0.016$ |
| Intrasession NTA map Similarity [r] | $0.534 \pm 0.031$ | $0.778 \pm 0.020$ |
| Intraindividual distance [mm] | $31.265 \pm 1.934$ | $13.717 \pm 1.129$ |

Similarly, Figure 2C and Figure 2D depict the individual NTA map and optimal target obtained from a 14-minute scan. As shown in the left column of Figure 2C and the left column of Figure 2D, the NTA map varied among individuals in the same scan, and optimal scalp sites were separated among individuals, which is consistent with previous studies (Ning et al., 2018; Cash et al., 2021). In the top row of Figure 2C and the top row of Figure 2D, target sites were constant across individuals over separate days, but variable among individuals across separate days in the bottom row of Figure 2C (i.e., Sub ID 189450), indicating the need for a stably personalized technique. The intrasession NTA map similarity for MDD and schizophrenia with AVH were 0.534 and 0.778, respectively.

When comparing individual target stability using 1 cm as the criterion, we found that the intraindividual distance was over 1 cm in both targeting MDD pathological network and targeting schizophrenia with AVH pathological network (Table 1). Additionally, we found that the stability was divided into two groups: stable group (top row of Figure 2C and Figure 2D) and unstable group (bottom row of Figure 2C and Figure 2D). The statistics of the two groups showed that 102 people were in the unstable group for targeting MDD pathological network, accounting for 76%; 32 people were in the stable group for targeting MDD pathological network, accounting for 24%; 72 people were in the unstable group for targeting schizophrenia with AVH pathological network, accounting for 54%; 62 people were in the stable targeting schizophrenia with AVH pathological network, accounting for 46%.

### 3.2. Increase the stability of individual sites by extending rsfMRI scan duration

We investigated the similarity of intrasession stimulation networks as scanning time increased. At a scanning time of 28 minutes, we observed that the intrasession stimulation network similarity at F3 was 0.750, while the intrasession stimulation network similarity at F3 was 0.779 (Supplementary Figure 3). We also calculated the intrasession stimulation network for all points in the search space and found a consistent increasing pattern, similar to that of a single target. In individuals targeting MDD pathological network, the intrasession stimulation network increased from 0.553 to 0.740 as the scan duration increased from 7 to 28 minutes (Figure 3A). Similarly, in individuals targeting schizophrenia with AVH pathological network, the intrasession stimulation network increased from 0.531 to 0.742 as the scan duration increased from 7 to 28 minutes (Figure 3B). These results indicate that retest reliability improves with longer scanning time.

**Figure 3.**
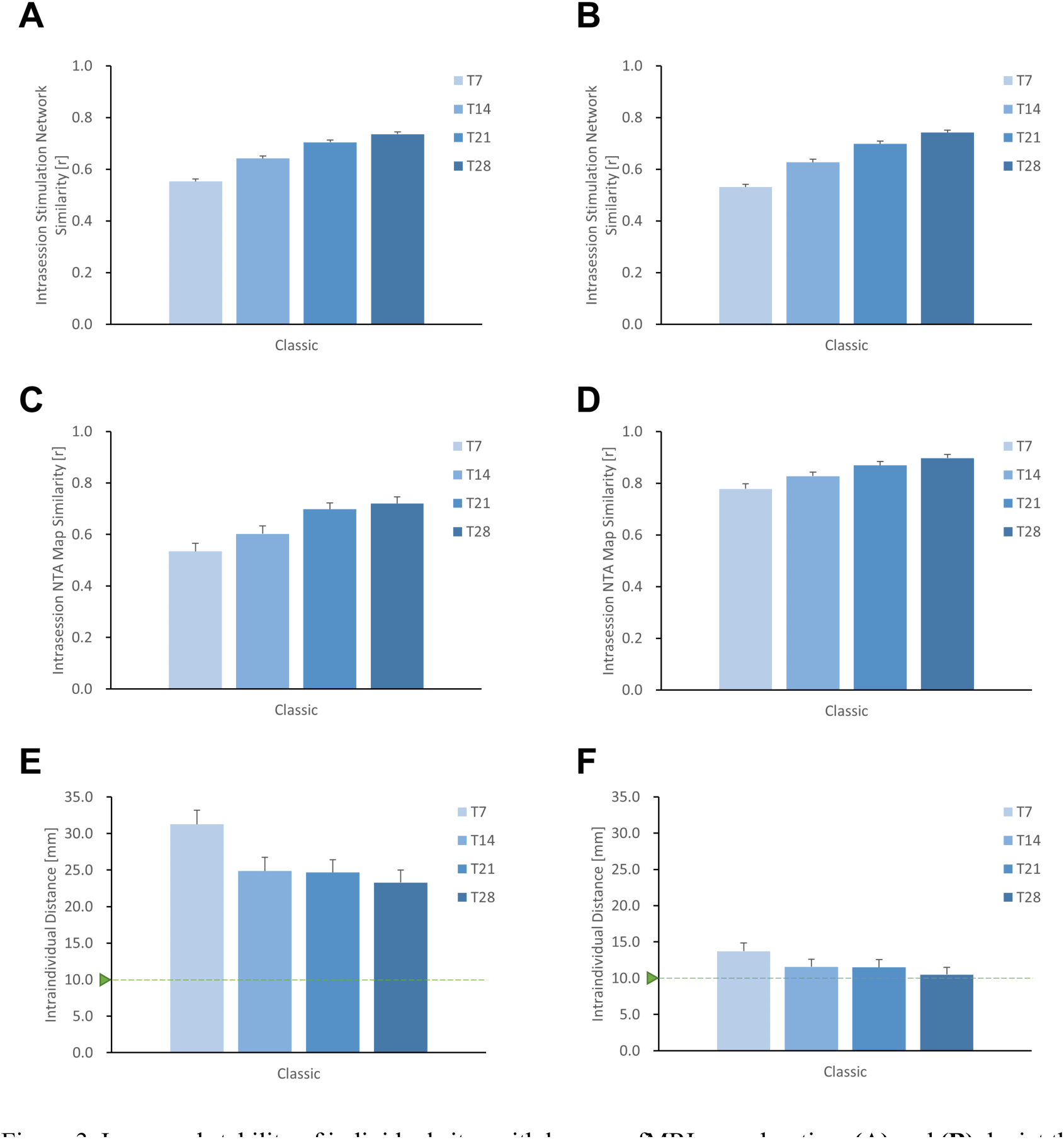
Improved stability of individual sites with longer rsfMRI scan duration. **(A)** and **(B)** depict the search space for MDD and schizophrenia with AVH respectively, showing that the stability of the stimulation network gradually increases with longer scanning time. Similarly, for the entire search space of **(C)** MDD and **(D)** AVH, the similarity of the NTA map also increases with extended scanning time. Additionally, when using the Classic method for optimizing the NTA map targeting either the **(E)** MDD network or the **(F)** AVH network, it is evident that the intraindividual distance of the target decreases with longer scanning time, although it remains higher than 1 cm.

Furthermore, the extension of scanning duration resulted in improved similarity of intrasession NTA maps (Figure 3C and 3D). In individuals targeting MDD pathological network, the correlation coefficient of the NTA map increased from 0.534 to 0.720 as the scan duration increased from 7 to 28 minutes. Similarly, in individuals targeting schizophrenia with AVH pathological network, the correlation coefficient of the NTA map increased from 0.778 to 0.897 with the same increase in scan duration. Both findings suggest that longer scan time enhances the reliability of the NTA map.

By using the classic method for optimization, we observed a gradual reduction in the distance of the optimal target within the individual with longer scanning time (Figure 3E and 3F). In individuals targeting MDD pathological network, the intraindividual distance decreased from 31.26 mm to 23.29 mm as the scan duration increased from 7 to 28 minutes. Similarly, in individuals targeting schizophrenia with AVH pathological network, the intraindividual distance decreased from 13.72 mm to 10.50 mm with the same increase in scan duration. However, both distances were still higher than 1cm. While longer scans lead to a decrease in intraindividual distance, additional searching methods are required to improve target stability.

### 3.3. Increase the stability of individual sites with cluster method

We evaluated the intraindividual distance of optimal sites (Figure 4) using a combination of the cluster method and 28-minute acquisition duration. The intraindividual distances were less than 1 cm both in individuals targeting MDD pathological network (9.23 ± 0.80 mm) and targeting schizophrenia with AVH pathological network (4.76 ± 0.40 mm), as depicted in Figures 4B and 4C. By using the cluster method, the intraindividual distances were reduced by 14 mm in MDD and 6 mm in schizophrenia with AVH, compared to the classic method and 28-minute acquisition duration. Although a longer acquisition duration results in a stable target, a shorter acquisition duration would be more practical. Figure 4B indicates that a 21-minute acquisition duration is the turning point for the stability of optimal targets.

**Figure 4.**
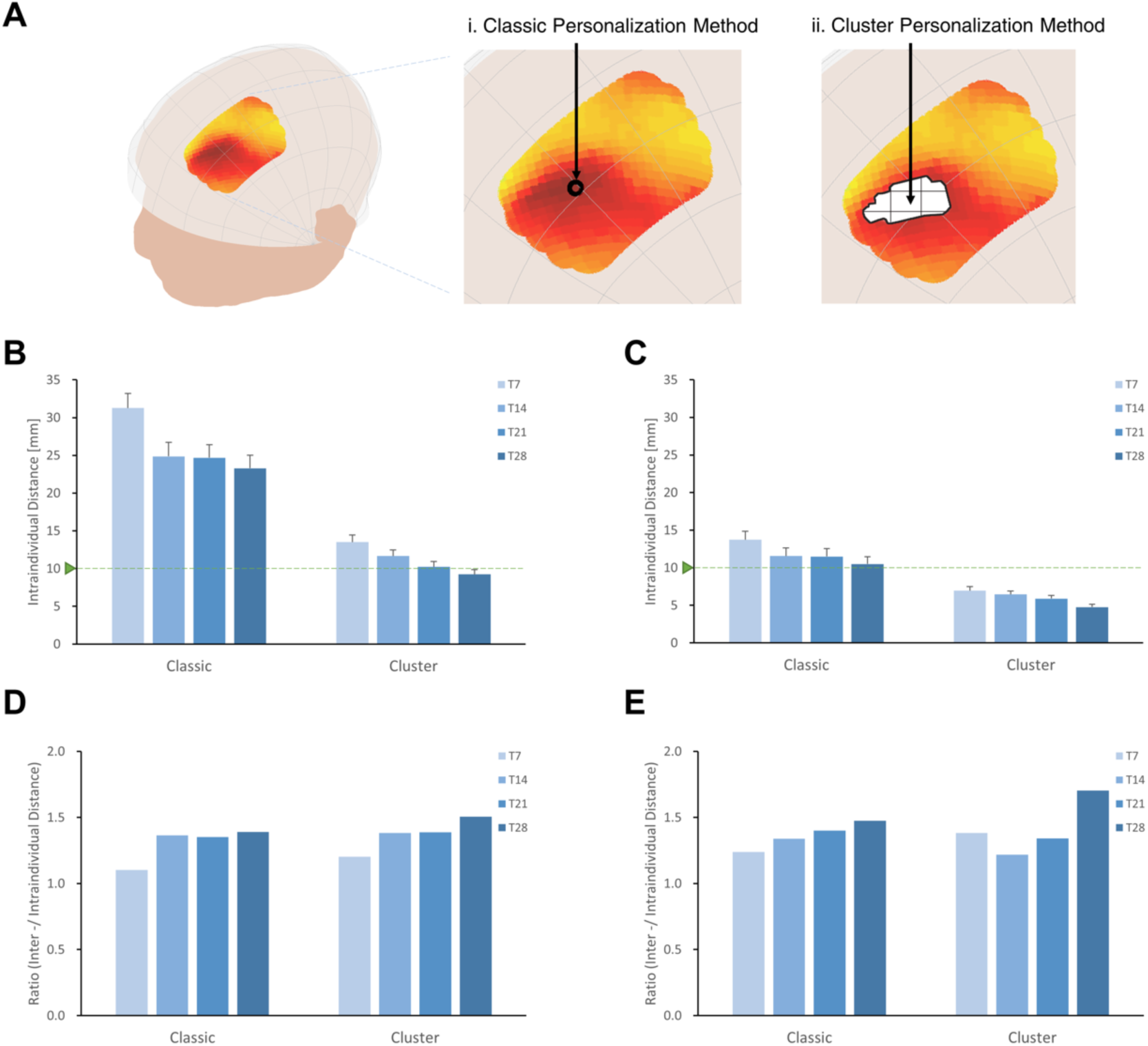
Increase the stability of individual sites with cluster method. **(A)** Classic method and Cluster method. Intraindividual distances between personalized targets were displayed for different methodologies and acquisition times (T7, T14, T21, T28), shown in targeting MDD pathological network **(B)** and schizophrenia with AVH pathological network **(C)**. Overall, the intraindividual distances with classic method were greater than those achieved with the cluster method when targeting MDD and schizophrenia with AVH pathological networks. In both MDD and schizophrenia with AVH, the intraindividual distances using the cluster method and T28 were found to be smaller than the spatial sensitivity of TMS (Deng et al., 2013). Furthermore, the cluster method and T28 exhibited the highest interindividual-to-intraindividual distance ratios both in MDD **(D)** and schizophrenia with AVH **(E)**.

We assessed the interindividual distance of optimal sites by employing the cluster method and a 28-minute acquisition duration (Supplementary Figure 4). In the case of targeting MDD pathological network, the interindividual distance was 13.89 mm, while for targeting schizophrenia with AVH pathological network, it was 8.11 mm. Utilizing the center-of-gravity calculation approach not only reduced intraindividual variance but also decreased interindividual variance, resulting in a 50% reduction in the interindividual distance compared to the classic method.

The cluster method proved to be more effective in identifying stable individual scalp sites for both targeting MDD and schizophrenia with AVH pathological networks when a 28-minute acquisition time was employed, as demonstrated by higher interindividual-to-intraindividual distance ratios (Figures 4C and D). Furthermore, even when the threshold was adjusted from 0.5% to 70%, the ratios remained consistent (Supplementary Figure 5). Specifically, the ratio was 1.51 for MDD and 1.70 for schizophrenia with AVH.

## 4. Discussion

In 1995, George et al. (George et al., 1995) pioneered the use of rTMS to modulate the DLPFC in patients with depression, resulting in successful alleviation of depressive symptoms through stimulation of DLPFC activity using 10Hz TMS. However, despite nearly three decades of evidence-based medical research, the average response rate for depression treatment remains around 50% (George et al., 2010; Brunoni et al., 2017; Miron et al., 2021), indicating the substantial potential for improving treatment efficacy. Personalized transcranial magnetic stimulation (TMS) interventions play a crucial role in enhancing treatment outcomes for psychiatric disorders. Among the personalized TMS interventions, connectivity-based targeting methods have emerged as a promising strategy (Fox et al., 2012; Cash et al., 2020). These methods leverage an individual’s functional connectivity patterns to target aberrant neural circuits and guide the selection of stimulation targets. A key aspect of achieving precision in TMS interventions lies in establishing a targeted pathway through functional connectivity to modulate the pathological networks associated with these disorders (Fox et al., 2014). In our previous study (Cao et al., 2023), we developed a network targeting model that enables precise and individualized interventions by specifically targeting the pathological networks based on the individual’s head surface. In the model, we employed group-level functional connectivity to construct the targeted pathway.

Utilizing individual functional connectivity has been shown to be valuable for enhancing clinical treatment prediction (Cash et al., 2020; Siddiqi et al., 2021; Kong et al., 2022). However, it is crucial to acknowledge the variability observed in individual-level functional connectivity, which can arise from both desirable and undesirable factors. Desirable variations reflect significant inter-individual differences in network organization and connectivity strength (Finn et al., 2015; Gratton et al., 2018), providing valuable insights for personalized TMS interventions. Conversely, undesirable variations stem from technical factors, measurement noise, or methodological limitations, introducing inconsistencies in the targeting process (Birn, 2012; Birn et al., 2013; Bright and Murphy, 2015; Ning et al., 2018; Cash et al., 2021). While desirable variations contribute to improving treatment efficacy, the undesirable variations resulting from technical factors can lead to instability in the personalized connectivity-based network targeting model. For instance, the intraindividual distance serves as a key index for evaluating the feasibility of a personalized model (Ning et al., 2018; Cash et al., 2021; Zhao et al., 2022). The utilization of individual rsFC for optimal TMS coil placement was found to be unstable due to the lower signal-to-noise ratio of rsfMRI (Dubois and Adolphs, 2016). In line with the personalized SGC-FC model (Ning et al., 2018), the application of the classic method in MDD with the personalized NTA model resulted in an intraindividual distance exceeding 30 mm, highlighting the challenge of achieving personalization.

To address the variability arising from technical factors and establish a stable personalized NTA model, we employed two strategies: extending the scan duration for data acquisition and identifying optimal stimulation sites at the computational level. The duration of rs-fMRI acquisition plays a critical role in determining a stable optimal site (Ning et al., 2018; Cash et al., 2021; Sun et al., 2022). We assessed the intraindividual distance using four scan durations in MDD and schizophrenia of AVH. In the classic method, the 28-minute acquisition time exhibited approximately 25% more reliability than the 7-minute acquisition time in MDD, as measured by the intraindividual distance index, with a 23% improvement observed in AVH. Considering that the optimal target may be an outlier singular point affecting the model’s stability, the cluster method utilized the similarity of NTA maps to identify reliable scalp sites by determining the center-of-gravity of the largest cluster, rather than relying solely on peak values (Cash et al., 2021; Zhao et al., 2022). Our findings demonstrated that the intraindividual distance using the cluster method and a 28-minute acquisition time was less than 1 cm, within the spatial sensitivity range of TMS (Deng et al., 2013), providing evidence of the effectiveness of the personalized NTA model. These strategies have partially addressed intra-individual variability.

Although capturing stable individual-specific functional network features may require several hours of scan data (Finn et al., 2015; Gratton et al., 2018), such an approach is impractical for therapeutic use. It is necessary to consider the tradeoff between scanning duration and the stability of individual targets (Sun et al., 2022). While a 28-minute scan is capable of acquiring a stable individual target, the turning point for stability was found to be a 21-minute scan, which is more practical. Additionally, alternative approaches for reliable functional connectivity estimation exist (Noble et al., 2019), including the use of a 7T magnetic field (Ning et al., 2018; Nemani and Lowe, 2021) and multivariate estimates of functional connectivity (Yoo et al., 2019). These strategies may be evaluated in future iterations of the personalized NTA model.

Furthermore, although strategies are helpful in decreasing the variance of undesirable technical factors, minimizing intra-individual variability may lead to the loss of desirable variations, as indicated by a decrease in interindividual variations (Supplementary Figure 4). The preservation of meaningful variations within individuals (Finn et al., 2015) and the establishment of a reasonable interindividual distance are crucial considerations. In an extreme example, when the interindividual distance is minimal, the best sites for all patients converge on one location. However, personalization becomes unnecessary if this one-site-fits-all method is successful. Clinical trials using neuronavigation targeting on a single site reported a response rate of approximately 50% (Blumberger et al., 2018; Li et al., 2020), indicating that the optimal sites for all patients were dispersed. However, the criteria for the interindividual distance between these optimal sites remain uncertain. Using the personalized NTA model, we found that the interindividual distance in our analysis (13.89 mm) was similar to the previous study (14.11 mm) (Cash et al., 2021) and close to the location variance of responders (19.45 mm), as demonstrated in Supplementary Figure 6.

Despite we detected reasonable interindividual variance in our study, the feasibility of implementing a personalized NTA model also relies on whether the variance among individuals exceeds the variance induced by different fMRI scans. Consistent with the findings of a prior study (Cash et al., 2021), we observed that the interindividual variance outweighed the intraindividual variation in both MDD and schizophrenia with AVH, as indicated by ratios greater than one. Furthermore, employing the cluster approach alongside a 28-minute scan duration yielded higher ratios compared to other combinations. The ratio index demonstrates the ability of the model to balance desirable and undesirable variations.

We incorporated individual functional connectivity into the NTA model to enhance the integration of the stimulation network and closely align it with the individual TMS effect (Cocchi and Zalesky, 2018a; Vink et al., 2018; Cash et al., 2020; Zhong et al., 2021; Klooster et al., 2022; Oliver et al., 2022). However, it is important to recognize that the variance in TMS treatment may not solely be attributed to the variance in the stimulation network but also to the variance in the pathological network. In our NTA model, we utilized a group-level coordinate-based meta-analysis as the pathological network (Cao et al., 2023). However, the pathological network could exist at the subgroup level (Drysdale et al., 2017; Cocchi and Zalesky, 2018b; Siddiqi et al., 2020) or at the individual-level (Gratton et al., 2020; Lynch et al., 2022). In future iterations of the personalized NTA model, it is crucial to consider psychosis biotypes (Drysdale et al., 2017; Siddiqi et al., 2020) and individual biomarkers (Gratton et al., 2020; Nawaz et al., 2021; Peng et al., 2023). Although personalized targeting has been successfully implemented in memory enhancement (Cash et al., 2022), it remains absent in the treatment of psychiatric disorders. The current study aims to bridge this gap by developing a feasible personalized NTA model for MDD and schizophrenia with AVH. It should be noted that our study has some limitations that need to be addressed through clinical experiments for validation. While a robust individualized NTA model set the path for accurate neuromodulation, it was insufficient to guide the precise positioning of TMS coils in clinical trials. Retrospective validation is essential before applicating personalized NTA model. Additionally, further research and exploration are required to uncover potential limitations and explore other avenues.

## 5. Conclusion

This study investigates the stability of a personalized connectivity-based network targeting model using the HCP dataset. Given the potential of individual functional connectivity to enhance the efficacy TMS, it is crucial to address technical variance of individual functional connectivity. To address this, we employed two strategies: extending the rsfMRI scan duration and utilizing an optimal target searching method. Our findings indicate that increasing the scan duration significantly improves the stability of the model. Furthermore, when combining the cluster method for identifying the optimal target with a longer scan duration, the intraindividual distance falls below the spatial resolution of TMS both in both MDD and schizophrenia with AVH. Although retrospective validation is necessary, our current model offers a feasible approach to obtaining stable personalized TMS targets for the treatment of psychiatric disorders.

## Author Contributions

**Zhengcao Cao**: Conceptualization, Formal analysis, Methodology, Investigation, Visualization, Data Curation, Software, Writing - original draft. **Xiang Xiao**: Conceptualization, Investigation, Supervision, Writing - review & editing. **Cong Xie**: Software. **Lijiang Wei**: Investigation. **Yihong Yang**: Funding acquisition, Conceptualization, Supervision, Writing - review & editing. **Chaozhe Zhu**: Funding acquisition, Conceptualization, Supervision, Writing - review & editing.

## Acknowledgments

The authors thank Zeqing Zheng and Farui Liu for their support during the study.

## Data Availability Statement

The participants, including the structure and resting-state functional MRI, are from the Human Connectome Project (HCP) database (https://db.humanconnectome.org/) and is openly available. The list of analyzed participants can be obtained upon request from C.Z.

## References

1. Ashburner, J. (2007). A fast diffeomorphic image registration algorithm. Neuroimage 38, 95–113. doi: 10.1016/j.neuroimage.2007.07.007.

2. Balderston, N. L., Roberts, C., Beydler, E. M., Deng, Z. De, Radman, T., Luber, B., et al. (2020). A generalized workflow for conducting electric field–optimized, fMRI-guided, transcranial magnetic stimulation. Nat. Protoc. 15, 3595–3614. doi: 10.1038/s41596-020-0387-4.

3. Barker, A. T., Jalinous, R., and Freeston, I. L. (1985). NON-INVASIVE MAGNETIC STIMULATION OF HUMAN MOTOR CORTEX. Lancet 325, 1106–1107. doi: 10.1016/S0140-6736(85)92413-4.

4. Birn, R. M. (2012). The role of physiological noise in resting-state functional connectivity. Neuroimage 62, 864–870. doi: 10.1016/j.neuroimage.2012.01.016.

5. Birn, R. M., Molloy, E. K., Patriat, R., Parker, T., Meier, T. B., Kirk, G. R., et al. (2013). The effect of scan length on the reliability of resting-state fMRI connectivity estimates. Neuroimage 83, 550–558. doi: 10.1016/j.neuroimage.2013.05.099.

6. Blumberger, D. M., Vila-rodriguez, F., Thorpe, K. E., Feffer, K., Noda, Y., Giacobbe, P., et al. (2018). Articles Effectiveness of theta burst versus high-frequency repetitive transcranial magnetic stimulation in patients with depression (THREE-D): a randomised non-inferiority trial. Lancet 391, 1683–1692. doi: 10.1016/S0140-6736(18)30295-2.

7. Bright, M. G., and Murphy, K. (2015). Is fMRI “noise” really noise? Resting state nuisance regressors remove variance with network structure. Neuroimage 114, 158–169. doi: 10.1016/j.neuroimage.2015.03.070.

8. Brunoni, A. R., Chaimani, A., Moffa, A. H., Razza, L. B., Gattaz, W. F., Daskalakis, Z. J., et al. (2017). Repetitive transcranial magnetic stimulation for the acute treatment of major depressive episodes a systematic review with network meta-analysis. JAMA Psychiatry 74, 143–152. doi: 10.1001/jamapsychiatry.2016.3644.

9. Cao, Z., Xiao, X., Zhao, Y., Jiang, Y., Xie, C., Paillère-Martinot, M. L., et al. (2023). Targeting the pathological network: Feasibility of network-based optimization of transcranial magnetic stimulation coil placement for treatment of psychiatric disorders. Front. Neurosci. 16, 1–15. doi: 10.3389/fnins.2022.1079078.

10. Cash, R. F. H., Cocchi, L., Lv, J., Wu, Y., Fitzgerald, P. B., and Zalesky, A. (2021). Personalized connectivity-guided DLPFC-TMS for depression: Advancing computational feasibility, precision and reproducibility. Hum. Brain Mapp. 42, 4155–4172. doi: 10.1002/hbm.25330.

11. Cash, R. F. H., Hendrikse, J., Fernando, K. B., Thompson, S., Suo, C., Fornito, A., et al. (2022). Personalized brain stimulation of memory networks. Brain Stimul. 15, 1300–1304. doi: 10.1016/j.brs.2022.09.004.

12. Cash, R. F. H., Weigand, A., Zalesky, A., Siddiqi, S. H., Downar, J., Fitzgerald, P. B., et al. (2020). Using Brain Imaging to Improve Spatial Targeting of Transcranial Magnetic Stimulation for Depression. Biol. Psychiatry 2, 1–8. doi: 10.1016/j.biopsych.2020.05.033.

13. Cash, R. F. H., Zalesky, A., Thomson, R. H., Tian, Y., Cocchi, L., and Fitzgerald, P. B. (2019). Subgenual Functional Connectivity Predicts Antidepressant Treatment Response to Transcranial Magnetic Stimulation: Independent Validation and Evaluation of Personalization. Biol. Psychiatry 86, e5–e7. doi: 10.1016/j.biopsych.2018.12.002.

14. Chou, P. H., Lin, Y. F., Lu, M. K., Chang, H. A., Chu, C. S., Chang, W. H., et al. (2021). Personalization of repetitive transcranial magnetic stimulation for the treatment of major depressive disorder according to the existing psychiatric comorbidity. Clin. Psychopharmacol. Neurosci. 19, 190–205. doi: 10.9758/cpn.2021.19.2.190.

15. Cocchi, L., and Zalesky, A. (2018a). Personalized Transcranial Magnetic Stimulation in Psychiatry. Biol. Psychiatry Cogn. Neurosci. Neuroimaging 3, 731–741. doi: 10.1016/j.bpsc.2018.01.008.

16. Cocchi, L., and Zalesky, A. (2018b). Personalized Transcranial Magnetic Stimulation in Psychiatry. Biol. Psychiatry Cogn. Neurosci. Neuroimaging 3, 731–741. doi: 10.1016/j.bpsc.2018.01.008.

17. Deng, Z. De, Lisanby, S. H., and Peterchev, A. V. (2013). Electric field depth-focality tradeoff in transcranial magnetic stimulation: Simulation comparison of 50 coil designs. Brain Stimul. 6, 1–13. doi: 10.1016/j.brs.2012.02.005.

18. Doucet, G. E., Lee, W. H., and Frangou, S. (2019). Evaluation of the spatial variability in the major resting-state networks across human brain functional atlases. Hum. Brain Mapp. 40, 4577–4587. doi: 10.1002/hbm.24722.

19. Drysdale, A. T., Grosenick, L., Downar, J., Dunlop, K., Mansouri, F., Meng, Y., et al. (2017). Resting-state connectivity biomarkers define neurophysiological subtypes of depression. Nat. Med. 23, 28–38. doi: 10.1038/nm.4246.

20. Du, R., Zhou, Q., Hu, T., Sun, J., Hua, Q., Wang, Y., et al. (2022). A landmark-based approach to locate symptom-specific transcranial magnetic stimulation targets of depression. Front. Psychol. 13, 1–9. doi: 10.3389/fpsyg.2022.919944.

21. Dubois, J., and Adolphs, R. (2016). Building a Science of Individual Differences from fMRI. Trends Cogn. Sci. 20, 425–443. doi: 10.1016/j.tics.2016.03.014.

22. Finn, E. S., Shen, X., Scheinost, D., Rosenberg, M. D., Huang, J., Chun, M. M., et al. (2015). Functional connectome fingerprinting : identifying individuals using patterns of brain connectivity. Nat. Publ. Gr. 18. doi: 10.1038/nn.4135.

23. Fischl, B., Rajendran, N., Busa, E., Augustinack, J., Hinds, O., Yeo, B. T. T., et al. (2008). Cortical folding patterns and predicting cytoarchitecture. Cereb. Cortex 18, 1973–1980. doi: 10.1093/cercor/bhm225.

24. Fitzgerald, P., Brown, T., Marston, N., Daskalakis, Z., Castella, A. De, and Kulkarni, J. (2003). Transcranial magnetic stimulation in the treatment of depression during pregnancy. Arch. Gen. Psychiatry 60, 1002–1008. doi: 10.1001/archpsyc.60.9.1002.

25. Fox, M. D., Buckner, R. L., Liu, H., Mallar Chakravarty, M., Lozano, A. M., and Pascual-Leone, A. (2014). Resting-state networks link invasive and noninvasive brain stimulation across diverse psychiatric and neurological diseases. Proc. Natl. Acad. Sci. U. S. A. 111, E4367–E4375. doi: 10.1073/pnas.1405003111.

26. Fox, M. D., Buckner, R. L., White, M. P., Greicius, M. D., and Pascual-Leone, A. (2012). Efficacy of transcranial magnetic stimulation targets for depression is related to intrinsic functional connectivity with the subgenual cingulate. Biol. Psychiatry 72, 595–603. doi: 10.1016/j.biopsych.2012.04.028.

27. Fox, M. D., Liu, H., and Pascual-Leone, A. (2013). Identification of reproducible individualized targets for treatment of depression with TMS based on intrinsic connectivity. Neuroimage 66, 151–160. doi: 10.1016/j.neuroimage.2012.10.082.

28. George, M. S., Lisanby, S. H., Avery, D., McDonald, W. M., Durkalski, V., Pavlicova, M., et al. (2010). Daily left prefrontal transcranial magnetic stimulation therapy for major depressive disorder: A sham-controlled randomized trial. Arch. Gen. Psychiatry 67, 507–516. doi: 10.1001/archgenpsychiatry.2010.46.

29. George, M. S., Wassermann, E. M., Williams, W. A., Callahan, A., Ketter, T. A., Basser, P., et al. (1995). Daily repetitive transcranial magnetic stimulation (rTMS) improves mood in depression. Neuroreport 6, 1853–1856. Available at: https://doi.org/10.1097/00001756-199510020-00008.

30. Gratton, C., Kraus, B. T., Greene, D. J., Gordon, E. M., Laumann, T. O., Nelson, S. M., et al. (2020). Defining Individual-Specific Functional Neuroanatomy for Precision Psychiatry. Biol. Psychiatry 88, 28–39. doi: 10.1016/j.biopsych.2019.10.026.

31. Gratton, C., Laumann, T. O., Nielsen, A. N., Schlaggar, B. L., Dosenbach, N. U. F., Petersen, S. E., et al. (2018). Functional Brain Networks Are Dominated by Stable Group and Individual Factors, Not Cognitive or Daily Article Functional Brain Networks Are Dominated by Stable Group and Individual Factors, Not Cognitive or Daily Variation. Neuron, 439–452. doi: 10.1016/j.neuron.2018.03.035.

32. Gray, J. P., Müller, V. I., Eickhoff, S. B., and Fox, P. T. (2020). Multimodal abnormalities of brain structure and function in major depressive disorder: A meta-analysis of neuroimaging studies. Am. J. Psychiatry 177, 422–434. doi: 10.1176/appi.ajp.2019.19050560.

33. Harita, S., Momi, D., Mazza, F., and Griffiths, J. D. (2022). Mapping Inter-individual Functional Connectivity Variability in TMS Targets for Major Depressive Disorder. Front. Psychiatry 13, 1–14. doi: 10.3389/fpsyt.2022.902089.

34. Herwig, U., Satrapi, P., and Schönfeldt-Lecuona, C. (2003). Using the International 10-20 EEG System for Positioning of Transcranial Magnetic Stimulation. Brain Topogr. 16, 95–99. doi: 10.1023/B:BRAT.0000006333.93597.9d.

35. Hill, J., Dierker, D., Neil, J., Inder, T., Knutsen, A., Harwell, J., et al. (2010). A surface-based analysis of hemispheric asymmetries and folding of cerebral cortex in term-born human infants. J. Neurosci. 30, 2268–2276. doi: 10.1523/JNEUROSCI.4682-09.2010.

36. Hoffman, R. E., Hawkins, K. A., Gueorguieva, R., Boutros, N. N., Rachid, F., Carroll, K., et al. (2003). Transcranial magnetic stimulation of left temporoparietal cortex and medication-resistant auditory hallucinations. Arch. Gen. Psychiatry 60, 49–56. doi: 10.1001/archpsyc.60.1.49.

37. Hoffman, R. E., Wu, K., Pittman, B., Cahill, J. D., Hawkins, K. A., Fernandez, T., et al. (2013). Transcranial magnetic stimulation of wernicke’s and right homologous sites to curtail voices: A randomized trial. Biol. Psychiatry 73, 1008–1014. doi: 10.1016/j.biopsych.2013.01.016.

38. Jiang, Y., Du, B., Chen, Y., Wei, L., Cao, Z., Zong, Z., et al. (2022). A scalp-measurement based parameter space: Towards locating TMS coils in a clinically-friendly way. Brain Stimul. 15, 924–926. doi: 10.1016/j.brs.2022.06.001.

39. Klooster, D. C. W., Ferguson, M. A., Boon, P. A. J. M., and Baeken, C. (2022). Personalizing Repetitive Transcranial Magnetic Stimulation Parameters for Depression Treatment Using Multimodal Neuroimaging. Biol. Psychiatry Cogn. Neurosci. Neuroimaging 7, 536–545. doi: 10.1016/j.bpsc.2021.11.004.

40. Kong, G., Wei, L., Wang, J., Zhu, C., and Tang, Y. (2022). The therapeutic potential of personalized connectivity-guided transcranial magnetic stimulation target over group-average target for depression. Brain Stimul. 15, 1063–1064. doi: 10.1016/j.brs.2022.07.054.

41. Kühn, S., and Gallinat, J. (2012). Quantitative meta-analysis on state and trait aspects of auditory verbal hallucinations in schizophrenia. Schizophr. Bull. 38, 779–786. doi: 10.1093/schbul/sbq152.

42. Lefaucheur, J. P., Aleman, A., Baeken, C., Benninger, D. H., Brunelin, J., Di Lazzaro, V., et al. (2020). Evidence-based guidelines on the therapeutic use of repetitive transcranial magnetic stimulation (rTMS): An update (2014–2018). Clin. Neurophysiol. 131, 474–528. doi: 10.1016/j.clinph.2019.11.002.

43. Lefaucheur, J. P., André-Obadia, N., Antal, A., Ayache, S. S., Baeken, C., Benninger, D. H., et al. (2014). Evidence-based guidelines on the therapeutic use of repetitive transcranial magnetic stimulation (rTMS). Clin. Neurophysiol. 125, 2150–2206. doi: 10.1016/j.clinph.2014.05.021.

44. Li, C. T., Cheng, C. M., Chen, M. H., Juan, C. H., Tu, P. C., Bai, Y. M., et al. (2020). Antidepressant Efficacy of Prolonged Intermittent Theta Burst Stimulation Monotherapy for Recurrent Depression and Comparison of Methods for Coil Positioning: A Randomized, Double-Blind, Sham-Controlled Study. Biol. Psychiatry 87, 443–450. doi: 10.1016/j.biopsych.2019.07.031.

45. Liu, W., Wei, D., Chen, Q., Yang, W., Meng, J., Wu, G., et al. (2017). Longitudinal test-retest neuroimaging data from healthy young adults in southwest China. Sci. Data 4, 1–9. doi: 10.1038/sdata.2017.17.

46. Lynch, C. J., Elbau, I. G., Ng, T. H., Wolk, D., Zhu, S., Ayaz, A., et al. (2022). Automated optimization of TMS coil placement for personalized functional network engagement. Neuron 110, 3263–3277.e4. doi: 10.1016/j.neuron.2022.08.012.

47. Menardi, A., Ozdemir, R. A., Momi, D., Tadayon, E., Boucher, P., Vallesi, A., et al. (2022). Effect of group-based vs individualized stimulation site selection on reliability of network-targeted TMS. Neuroimage 264. doi: 10.1016/j.neuroimage.2022.119714.

48. Mira-Dominguez, O., Mills, B. D., Carpenter, S. D., Grant, K. A., Kroenke, C. D., Nigg, J. T., et al. (2014). Connectotyping: Model based fingerprinting of the functional connectome. PLoS One 9. doi: 10.1371/journal.pone.0111048.

49. Miron, J.-P., Jodoin, V. D., Lespérance, P., and Blumberger, D. M. (2021). Repetitive transcranial magnetic stimulation for major depressive disorder: basic principles and future directions. Ther. Adv. Psychopharmacol. 11, 204512532110426. doi: 10.1177/20451253211042696.

50. Mueller, S., Wang, D., Fox, M. D., Yeo, B. T. T., Sepulcre, J., Sabuncu, M. R., et al. (2013). Individual Variability in Functional Connectivity Architecture of the Human Brain. Neuron 77, 586–595. doi: 10.1016/j.neuron.2012.12.028.

51. Nawaz, U., Lee, I., Beermann, A., Eack, S., Keshavan, M., and Brady, R. (2021). Individual Variation in Functional Brain Network Topography is Linked to Schizophrenia Symptomatology. Schizophr. Bull. 47, 180–188. doi: 10.1093/schbul/sbaa088.

52. Nemani, A., and Lowe, M. J. (2021). Seed-based test–retest reliability of resting state functional magnetic resonance imaging at 3T and 7T. Med. Phys. 48, 5756–5764. doi: 10.1002/mp.15210.

53. Ning, L., Makris, N., Camprodon, J. A., and Rathi, Y. (2018). Brain Stimulation Limits and reproducibility of resting-state functional MRI de fi nition of DLPFC targets for neuromodulation. Brain Stimul., 1–10. doi: 10.1016/j.brs.2018.10.004.

54. Noble, S., Scheinost, D., and Constable, R. T. (2019). NeuroImage A decade of test-retest reliability of functional connectivity : A systematic review and meta-analysis. Neuroimage 203, 116157. doi: 10.1016/j.neuroimage.2019.116157.

55. Oliver, L. D., Hawco, C., Viviano, J. D., and Voineskos, A. N. (2022). From the Group to the Individual in Schizophrenia Spectrum Disorders: Biomarkers of Social Cognitive Impairments and Therapeutic Translation. Biol. Psychiatry 91, 699–708. doi: 10.1016/j.biopsych.2021.09.007.

56. Paillère-Martinot, M. L., Galinowski, A., Plaze, M., Andoh, J., Bartrés-Faz, D., Bellivier, F., et al. (2017). Active and placebo transcranial magnetic stimulation effects on external and internal auditory hallucinations of schizophrenia. Acta Psychiatr. Scand. 135, 228–238. doi: 10.1111/acps.12680.

57. Paillère Martinot, M. L., Galinowski, A., Ringuenet, D., Gallarda, T., Lefaucheur, J. P., Bellivier, F., et al. (2010). Influence of prefrontal target region on the efficacy of repetitive transcranial magnetic stimulation in patients with medication-resistant depression: A [18F]-fluorodeoxyglucose PET and MRI study. Int. J. Neuropsychopharmacol. 13, 45–59. doi: 10.1017/S146114570900008X.

58. Park, K. Y., Lee, J. J., Dierker, D., Marple, L. M., Hacker, C. D., Roland, J. L., et al. (2020). Mapping language function with task-based vs. resting-state functional MRI. PLoS One 15, 1–16. doi: 10.1371/journal.pone.0236423.

59. Peng, X., Liu, Q., Hubbard, C. S., Wang, D., Zhu, W., Fox, M. D., et al. (2023). Robust dynamic brain coactivation states estimated in individuals. Sci. Adv. 9. doi: 10.1126/sciadv.abq8566.

60. Rajkowska, G., and Goldman-Rakic, P. S. (1995). Cytoarchitectonic definition of prefrontal areas in normal human cortex: I. Remapping of areas 9 and 46 and relationship to the Talairach coordinate system. Cereb. Cortex 5, 307–322. Available at: https://doi.org/10.1093/cercor/5.4.307.

61. Rossini, P. M., Rossini, L., and Ferreri, F. (2010). Transcranial magnetic stimulation: A review. IEEE Eng. Med. Biol. Mag. 29, 84–95. doi: 10.1109/MEMB.2009.935474.

62. Sale, M. V., Mattingley, J. B., Zalesky, A., and Cocchi, L. (2015). Imaging human brain networks to improve the clinical efficacy of non-invasive brain stimulation. Neurosci. Biobehav. Rev. 57, 187–198. doi: 10.1016/j.neubiorev.2015.09.010.

63. Saturnino, G. B., Puonti, O., Nielsen, J. D., Antonenko, D., Madsen, K. H., and Thielscher, A. (2019). SimNIBS 2.1: A Comprehensive Pipeline for Individualized Electric Field Modelling for Transcranial Brain Stimulation. Brain Hum. Body Model., 3–25. doi: 10.1007/978-3-030-21293-3_1.

64. Siddiqi, S. H., Taylor, S. F., Cooke, D., Pascual-Leone, A., George, M. S., and Fox, M. D. (2020). Distinct symptom-specific treatment targets for circuit-based neuromodulation. Am. J. Psychiatry 177, 435–446. doi: 10.1176/appi.ajp.2019.19090915.

65. Siddiqi, S. H., Weigand, A., Pascual-Leone, A., and Fox, M. D. (2021). Identification of Personalized Transcranial Magnetic Stimulation Targets Based on Subgenual Cingulate Connectivity: An Independent Replication. Biol. Psychiatry, 1–2. doi: 10.1016/j.biopsych.2021.02.015.

66. Sun, J., Du, R., Zhang, B., Hua, Q., Wang, Y., Zhang, Y., et al. (2022). Minimal scanning duration for producing individualized repetitive transcranial magnetic stimulation targets. Brain Imaging Behav. 16, 2637–2646. doi: 10.1007/s11682-022-00720-y.

67. Thielscher, A., Antunes, A., and Saturnino, G. B. (2015). Field modeling for transcranial magnetic stimulation: A useful tool to understand the physiological effects of TMS? in 2015 37th Annual International Conference of the IEEE Engineering in Medicine and Biology Society (EMBC) (IEEE), 222–225. doi: 10.1109/EMBC.2015.7318340.

68. Thielscher, A., and Kammer, T. (2004). Electric field properties of two commercial figure-8 coils in TMS: Calculation of focality and efficiency. Clin. Neurophysiol. 115, 1697–1708. doi: 10.1016/j.clinph.2004.02.019.

69. Thomson, R. H., Cleve, T. J., Bailey, N. W., Rogasch, N. C., Maller, J. J., Daskalakis, Z. J., et al. (2013). Blood oxygenation changes modulated by coil orientation during prefrontal transcranial magnetic stimulation. Brain Stimul. 6, 576–581. doi: 10.1016/j.brs.2012.12.001.

70. Vila-Rodriguez, F., and Frangou, S. (2021). Individualized functional targeting for rTMS: A powerful idea whose time has come? Hum. Brain Mapp. 42, 4079– 4080. doi: 10.1002/hbm.25543.

71. Vink, J. J. T., Mandija, S., Petrov, P. I., van den Berg, C. A. T., Sommer, I. E. C., and Neggers, S. F. W. (2018). A novel concurrent TMS-fMRI method to reveal propagation patterns of prefrontal magnetic brain stimulation. Hum. Brain Mapp. 39, 4580–4592. doi: 10.1002/hbm.24307.

72. Wagner, T. A., Zahn, M., Grodzinsky, A. J., and Pascual-Leone, A. (2004). Three-dimensional head model simulation of transcranial magnetic stimulation. IEEE Trans. Biomed. Eng. 51, 1586–1598. doi: 10.1109/TBME.2004.827925.

73. Weigand, A., Horn, A., Caballero, R., Cooke, D., Stern, A. P., Taylor, S. F., et al. (2018). Prospective Validation That Subgenual Connectivity Predicts Antidepressant Efficacy of Transcranial Magnetic Stimulation Sites. Biol. Psychiatry 84, 28–37. doi: 10.1016/j.biopsych.2017.10.028.

74. Xiao, X., Yu, X., Zhang, Z., Zhao, Y., Jiang, Y., Li, Z., et al. (2018). Transcranial brain atlas. Sci. Adv. 4. doi: 10.1126/sciadv.aar6904.

75. Yan, C. G., Wang, X. Di, Zuo, X. N., and Zang, Y. F. (2016). DPABI: Data Processing & Analysis for (Resting-State) Brain Imaging. Neuroinformatics 14, 339–351. doi: 10.1007/s12021-016-9299-4.

76. Yoo, K., Rosenberg, M. D., Noble, S., Scheinost, D., Constable, R. T., and Chun, M. M. (2019). NeuroImage Multivariate approaches improve the reliability and validity of functional connectivity and prediction of individual behaviors. Neuroimage 197, 212–223. doi: 10.1016/j.neuroimage.2019.04.060.

77. Zhao, N., Yue, J., Feng, Z. J., Qiao, Y., Ge, Q., Yuan, L. X., et al. (2022). The Location Reliability of the Resting-State fMRI FC of Emotional Regions Towards rTMS Therapy. Neuroinformatics 20, 1055–1064. doi: 10.1007/s12021-022-09585-4.

78. Zhong, G., Yang, Z., and Jiang, T. (2021). Precise Modulation Strategies for Transcranial Magnetic Stimulation : Advances and Future Directions. Neurosci. Bull. 37, 1718–1734. doi: 10.1007/s12264-021-00781-x.

