## Supplementary Figure1 for "Personalized connectivity-based network targeting model of TMS for treatment of psychiatric disorders: computational feasibility and reproducibility"

*Supplementary Materials*

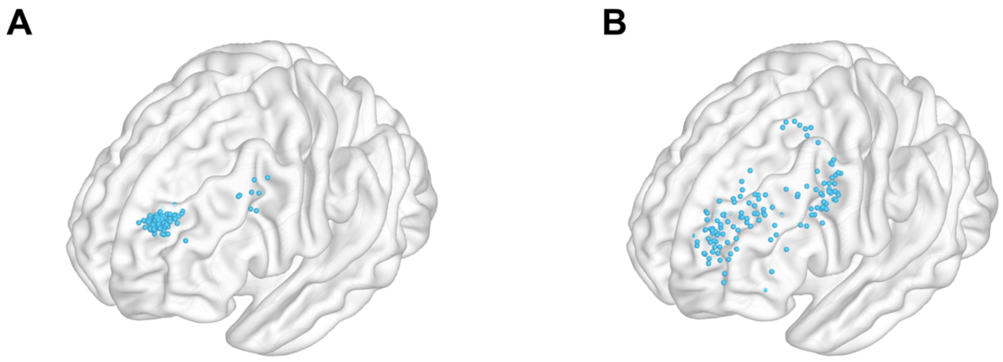


Supplementary Figure 1. The comparison between optimal targets with different models under 28-minutes scans. **(A)** Optimal targets with network targeting model using group-level rsFC. **(B)** Optimal targets with network targeting model using individual rsFC.

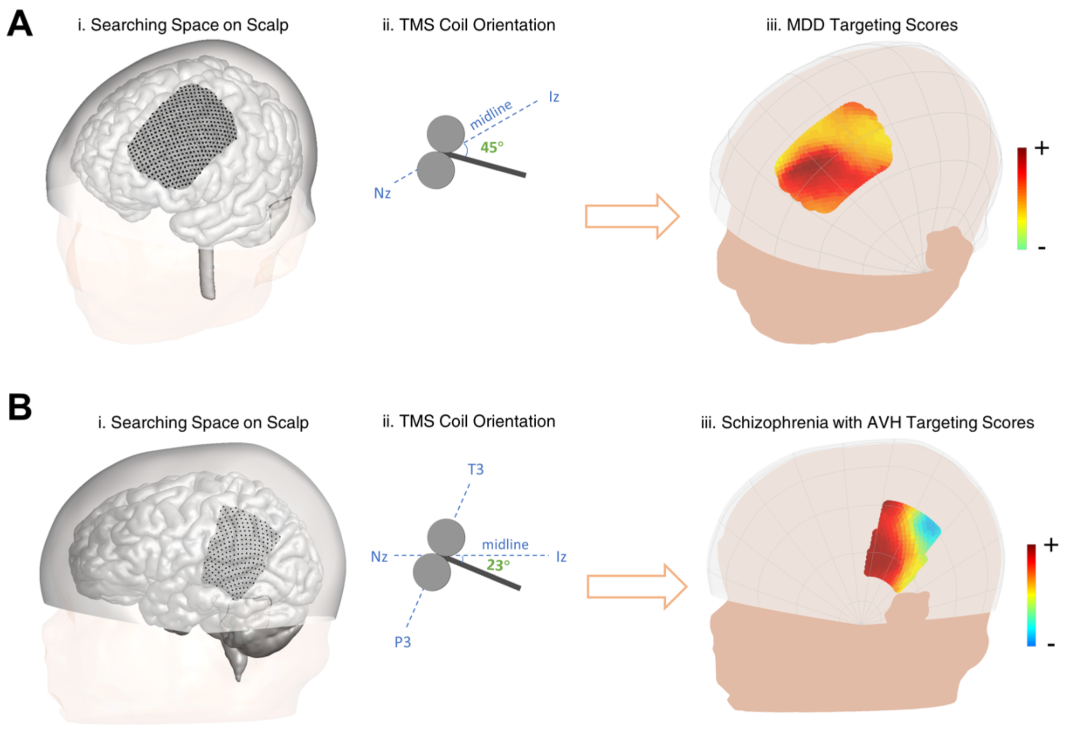


Supplementary Figure 2. Search space, coil orientation, and NTA map targeting MDD and schizophrenia with AVH pathological networks. **(A)** Illustration of a representative individual's search space for MDD treatment (i). The search space is depicted by 462 positions (black dots). A fixed coil orientation (45° from the midline, ii) is maintained across the positions. NTA values are computed for each position-orientation pair, resulting in the corresponding MDD NTA map displayed within the search space (iii). **(B)** Illustration of a representative individual's search space for schizophrenia with AVH treatment (i). The search space is represented by 246 positions (black dots). A fixed coil orientation (23° from the midline, ii) is applied uniformly across the positions. NTA values are calculated for each position-orientation combination, leading to the corresponding schizophrenia with AVH NTA map shown within the search space (iii).

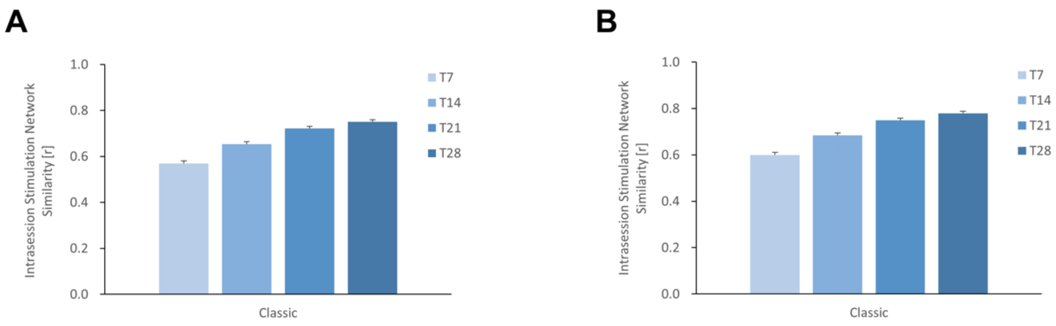


Supplementary Figure 3. Similarity of stimulation network across scan times for a specific scalp location. The figure demonstrates the calculation of similarity between stimulation networks acquired during different scanning durations. Both **(A)** F3 and **(B)** TP3 scalp locations show an increasing intrasession stimulation network similarity as the scanning time is extended.

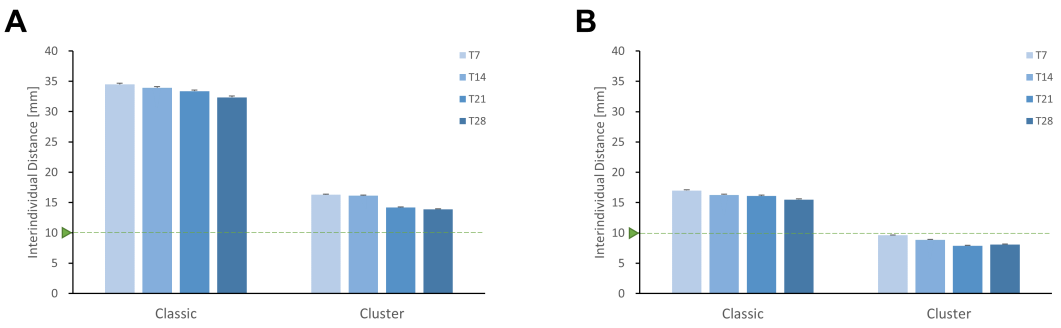


Supplementary Figure 4. Interindividual distances. In order to assess the consistency across individuals, the distances between optimal targets obtained from the same scan but different individuals were calculated using either the Classic or Cluster method. The results are presented for **(A)** targeting the MDD pathological network and **(B)** targeting schizophrenia with AVH pathological network.

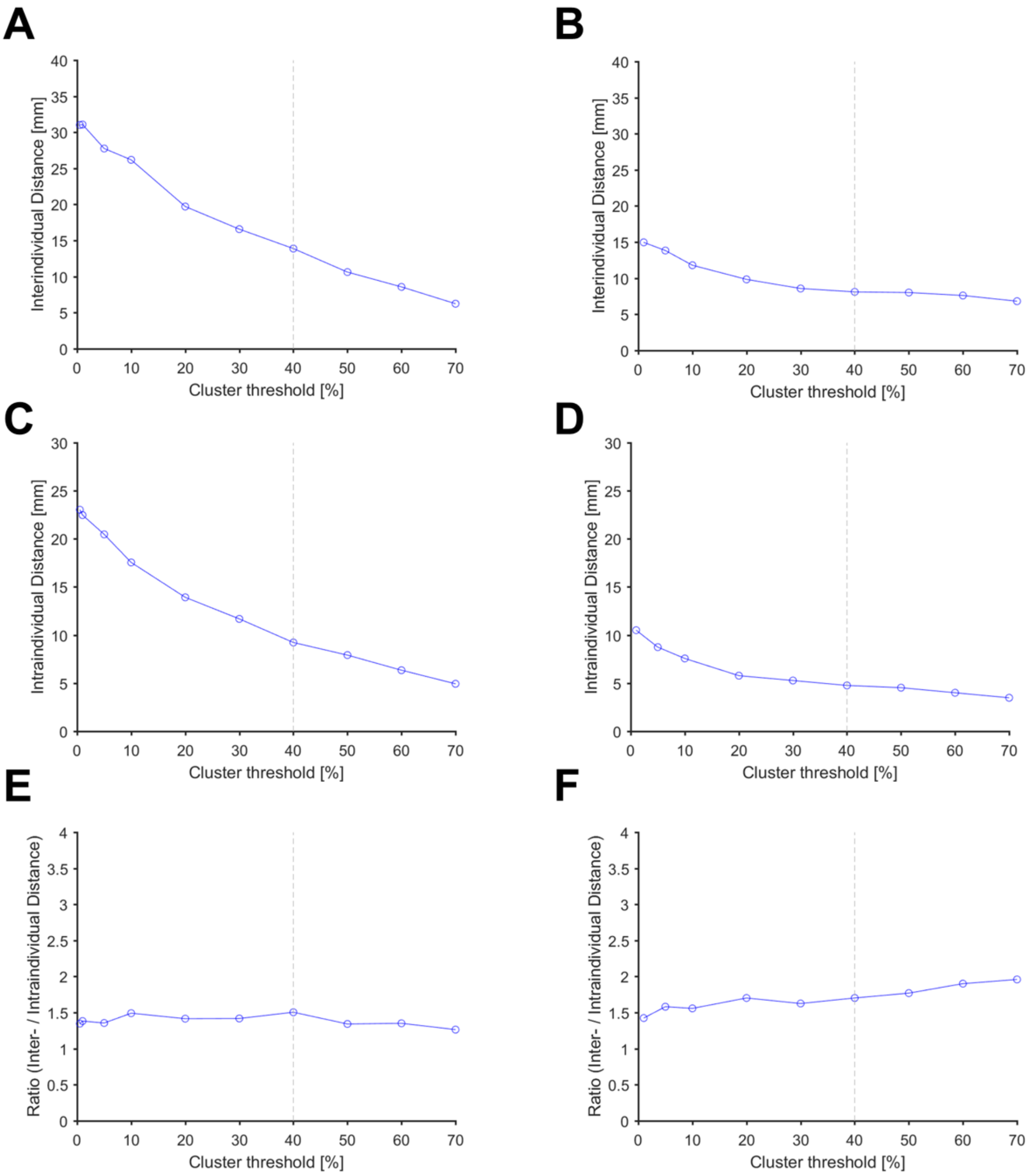


Supplementary Figure 5. The impact of cluster threshold on the results. The interindividual distances were measured when targeting the MDD pathological network **(A)** and the schizophrenia with AVH pathological network **(B)**. The intraindividual distances were measured when targeting the MDD pathological network **(C)** and the schizophrenia with AVH pathological network **(D)**. The ratios were measured when targeting the MDD pathological network **(E)** and the schizophrenia with AVH pathological network **(F)**. To determine the appropriate threshold, the ratios were examined for both the MDD pathological network and the schizophrenia with AVH pathological network. It was observed that the ratios remained consistent for both networks. Therefore, a threshold of 40% was selected based on the maximum ratio value observed in the MDD network. Additionally, the interindividual distance obtained with the 40% threshold (13.89 mm) was comparable to the distance (14.11 mm) reported in a previous study (Cash et al., 2021).

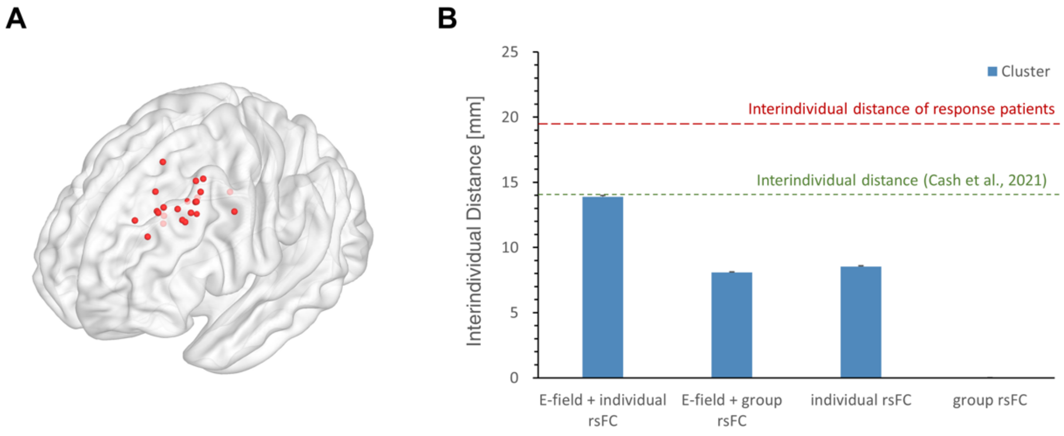


Supplementary Figure 6. Comparison of interindividual distances among different models. TMS treatment responders, defined as individuals with a ≥50% reduction in clinical scores, were selected from previous studies (Paillère Martinot et al., 2010; Weigand et al., 2018). The cortical targeting sites are illustrated in **(A)**. The interindividual distance between these targeting sites of responders was used to measure the location variance among individuals. **(B)** Our findings revealed that the variance of individual targeting sites obtained using the personalized NTA model was similar to that reported in the previous study (Cash et al., 2021), and it closely matched the location variance of the responders. Moreover, the variation observed when considering both individual E-field and individual rsFC was higher than when considering either individual E-field or individual rsFC alone. This suggests that personalized TMS treatment should account for both individual cortical characteristics and functional connectivity.
